# Aging and white matter microstructure and macrostructure: a longitudinal multi-site diffusion MRI study of 1,184 participants

**DOI:** 10.1101/2022.02.10.479977

**Authors:** Kurt G Schilling, Derek Archer, Fang-Cheng Yeh, Francois Rheault, Leon Y Cai, Colin Hansen, Qi Yang, Karthik Ramdass, Andrea Shafer, Susan Resnick, Kimberly R. Pechman, Katherine A. Gifford, Timothy J. Hohman, Angela Jefferson, Adam W Anderson, Hakmook Kang, Bennett A Landman

## Abstract

Quantifying the microstructural and macrostructural geometrical features of the human brain’s connections is necessary for understanding normal aging and disease. Here, we examine brain white matter diffusion magnetic resonance imaging data from one cross-sectional and two longitudinal datasets totaling in 1184 subjects and 2236 sessions of people aged 50-97 years. Data was drawn from well-established cohorts, including the Baltimore Longitudinal Study of Aging dataset, Cambridge Centre for Ageing Neuroscience dataset, and the Vanderbilt Memory & Aging Project. Quantifying 4 microstructural features and, for the first time, 11 macrostructure-based features of volume, area, and length across 120 white matter pathways, we apply linear mixed effect modeling to investigate changes in pathway-specific features over time, and document large age associations within white matter. Conventional diffusion tensor microstructure indices are the most age-sensitive measures, with positive age associations for diffusivities and negative age associations with anisotropies, with similar patterns observed across all pathways. Similarly, pathway shape measures also change with age, with negative age associations for most length, surface area, and volume-based features. A particularly novel finding of this study is that while trends were homogeneous throughout the brain for microstructure features, macrostructural features demonstrated heterogeneity across pathways, whereby several projection, thalamic, and commissural tracts exhibited more decline with age compared to association and limbic tracts. The findings from this large-scale study provide a comprehensive overview of the age-related decline in white matter and demonstrate that macrostructural features may be more sensitive to heterogeneous white matter decline. Therefore, leveraging macrostructural features may be useful for studying aging and could have widespread implications for a variety of neurodegenerative disorders.

## Introduction

To better understand changes related to normal aging, and differences due to disease, it is necessary to characterize how and where the brain changes with age. Studies using magnetic resonance imaging (MRI) have shown that the brain undergoes significant changes with age. Most studies focus on gray matter of the brain, where correlations between cortical volumes and age have been consistently described. These findings provide evidence of heterogenous patterns of normal age-related changes (3–8), with detectable differences in neurological diseases and disorders (9–13).

While white matter appears relatively homogenous on conventional structural MRI, diffusion MRI and subsequent fiber tractography enables investigation of individual fiber pathways of the brain. To date, most diffusion MRI studies of aging characterize features of tissue *microstructure* using cross-sectional datasets. For example, diffusion tensor imaging (DTI) shows fractional anisotropy (FA) is negatively associated with age, and mean diffusivity (MD) positively associated with age across white matter pathways (14–17), and have shown that advanced multicompartment diffusion modeling also provides sensitive measures of age-related microstructural changes (18–21). Microstructural features of these fiber pathways are biologically relevant in aging research as demyelination is thought to occur in a heterogeneous manner, whereby late-myelinating fiber pathways exhibit neurodegeneration prior to other fiber pathways. This idea, known as the myelodegeneration hypothesis, has recently been supported by a large-scale diffusion MRI study leveraging data from the UK Biobank (n=7,167) (22). Specifically, they found disproportional age-related differences in fiber pathways projecting to/from the prefrontal cortex.

While diffusion-based microstructure has been widely studied in aging, the *macrostructural* features of these fiber pathways play a pivotal role along the aging continuum; however, they have yet to be studied. As recently described(2), these macrostructural properties – descriptions of lengths, areas, and volumes - can be used to describe the geometrical and connectivity features of fiber bundles. The incorporation of these features into the study of aging and aging-related disorders could provide an additional avenue to elucidate the mechanisms driving white matter neurodegeneration. Given our prior knowledge, our hypothesis is that microstructural and macrostructural features will be disproportionately affected in fiber tracts projecting to/from the prefrontal cortex along the aging continuum.

To address our hypothesis, we will leverage three well-established cohorts of aging, including two longitudinal cohorts [Baltimore Longitudinal Study of Aging (BLSA)(23), Vanderbilt Memory & Aging Project (VMAP)(24)] and one cross-sectional cohort [Cambridge Centre for Ageing and Neuroscience (Cam-CAN)(25)]. Within these cohorts, automated tractography segmentation will be conducted within 120 white matter tracts, including association, limbic, projection (including thalamic and striatal), and commissural tracts. We will then quantify 4 microstructural and 11 macrostructural features within these tracts to determine if these metrics exhibit disproportionate age-related decline.

## Methods

### Data

This study used data from three datasets, summarized in **Table 1**, and contained a total of 1184 subjects and 2236 sessions of healthy subjects aged 50-97 years. All datasets were filtered to exclude subjects with diagnoses of mild cognitive impairment, Alzheimer’s disease, or dementia at baseline, or if they developed these conditions during the follow-up interval. Finally, datasets were filtered in order to focus on subjects aged 50+, due to limited samples sizes of each dataset with subjects below 50 years old.

**Table 1.** Datasets. This study used 3 longitudinal and cross-sectional datasets, with a total of 1184 subjects and 2236 sessions of healthy subjects aged 50-97 years.

| Dataset | Number Subjects | Number of Sessions | Age |
| --- | --- | --- | --- |
| Baltimore Longitudinal Study of Aging | 641<br>280 M | 1322<br>Range [1 8] | [50 97]<br>73.3 ± 9.8 |
| Cambridge Centre for Ageing Neuroscience | 356<br>179 M | 356<br>Range [1] | [50 88]<br>67.9 ± 10.3 |
| Vanderbilt Memory & Aging Project | 187<br>113 M | 558<br>Range [1 4] | [60 95]<br>74.2 ± 7.0 |
|  | **1184**<br>**572 M** | **2236**<br>**Range [1 8]** | **[50 97]**<br>**72.6 ± 9.5** |

First, was the Baltimore Longitudinal Study of Aging (BLSA) dataset, with 641 subjects scanned multiple times ranging from 1 and 8 sessions, and time between scans ranging from 1 to 10 years, yielding a total of 1322 diffusion datasets. Diffusion MRI data was acquired on a 3T Philips Achieva scanner (32 gradient directions, b-value=700s/mm2, TR/TE=7454/75ms, reconstructed voxel size=0.81×0.81×2.2mm, reconstruction matrix=320×320, acquisition matrix=115× 115, field of view=260×260mm). Second, was data from the Vanderbilt Memory & Aging Project (VMAP), with 187 subjects, scanned between 1-4 sessions, with a total of 558 diffusion datasets. Diffusion MRI data was acquired on a 3T Philips Achieva scanner (32 gradient directions, b-value=1000s/mm2, reconstructed voxel size=2×2×2mm). Third, was data from the Cambridge Centre for Ageing and Neuroscience (Cam-CAN) data repository (25) with 356 subjects, each scanned once using a 3T Siemens TIM Trio scanner with a 32-channel head coil (30 directions at b-value=1000s/mm2, 30 directions at b-value=2000s/mm2, reconstructed voxel size=2×2×2mm). All human datasets from Vanderbilt University were acquired after informed consent under supervision of the appropriate Institutional Review Board. All additional datasets are freely available and unrestricted for noncommercial research purposes. This study accessed only deidentified patient information.

### Processing

For every session, sets of white matter pathways were virtually dissected using two automated fiber tractography pipelines, TractSeg(1) and Automatic Track Recognition (ATR) (26). Two methods, based on different technological and anatomical principles of tractography segmentation were selected to emphasize generalizability of results across choices of different workflow(27).

Throughout the manuscript, TractSeg analysis is presented as primary results, and ATR as supplementary.

Briefly, TractSeg was based on convolutional neural networks and performed bundle-specific tractography based on a field of estimated fiber orientations (1). We implemented the dockerized version at (https://github.com/MIC-DKFZ/TractSeg), which generated fiber orientations using constrained spherical deconvolution with the MRtrix3 software (28). TractSeg resulted in 72 bundles, visualized in **Figure 1**, including association, limbic, commissural, thalamic, striatal, and projection and cerebellar pathways.

**Figure 1.**
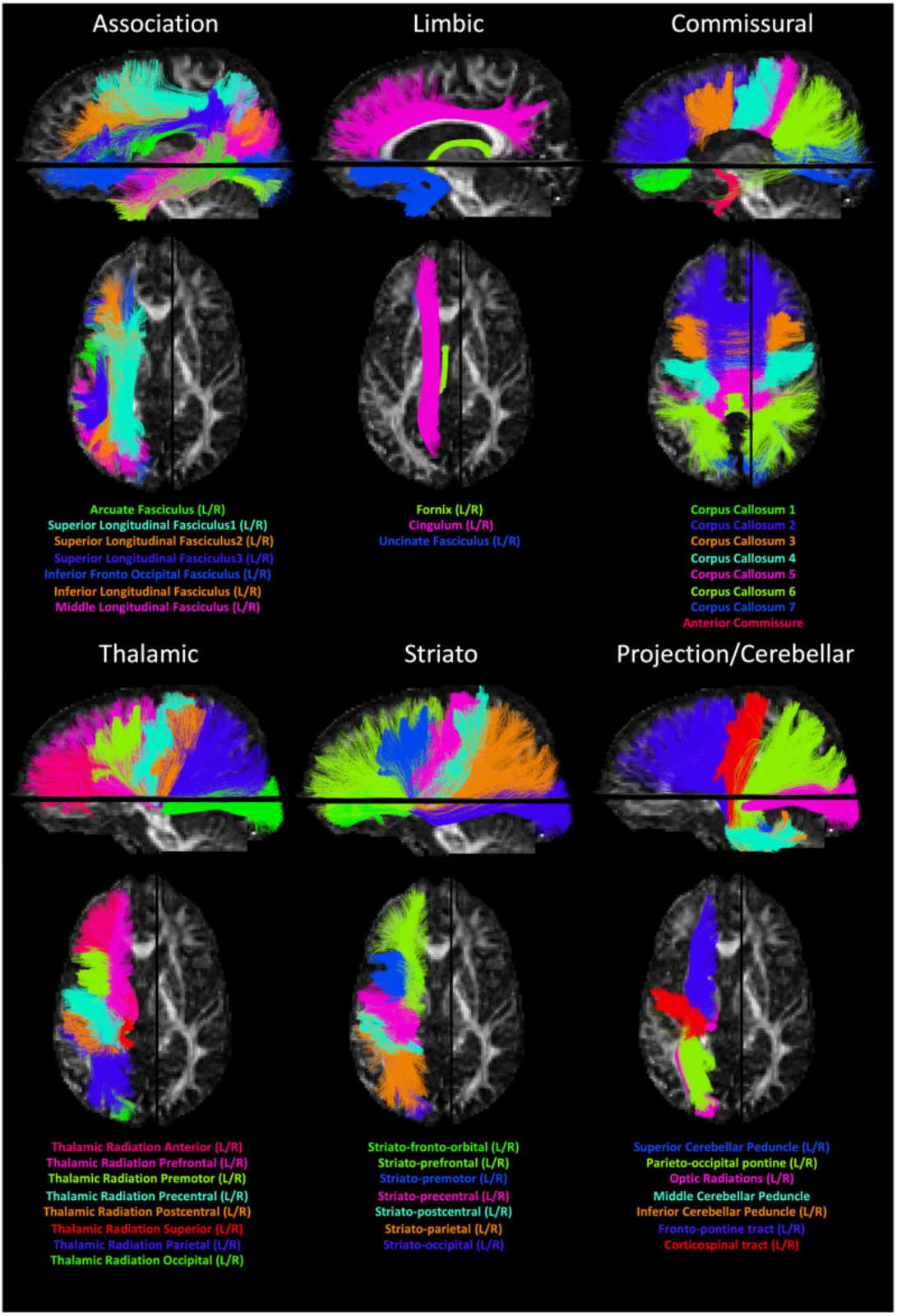
We investigated microstructure and macrostructure features of 71 pathways virtually dissected using TractSeg(1), visualized and organized into association, limbic, commissural, thalamic, striatal, and projection and cerebellar pathways.

ATR was performed in DSI Studio software using batch automated fiber tracking (26). Data were reconstructed using generalized q-sampling imaging(29) with a diffusion sampling length ratio of 1.25. A deterministic fiber tracking algorithm (30) was used in combination with anatomical priors from a tractography atlas (26) to map all pathways using inclusion and exclusion regions of interest. Topology-informed pruning (31) was applied to the tractography with 16 iterations to remove false connections. The Dockerized source code is available at http://dsi-studio.labsolver.org. ATR resulted. In 49 bundles, visualized in. **Supplementary Figure 1**, including association, limbic, commissural, thalamic, and projection pathways.

For each session, and every pathway, several features were extracted. Four microstructural features included DTI metrics of FA, MD, radial diffusivity (RD), and axial diffusivity (AD). The 11 macrostructure-based features extracted from each pathway (details described in (2)) are based on length (mean length, span, diameter; units of mm), area (total surface area, total area of end regions; units of mm^2^), volume (total volume, trunk volume, branch volume; units of mm^3^), and shape (curl, elongation, irregularity; unitless). Summary descriptions and equations for all macrostructural features are shown in **Table 2**.

**Table 2.**
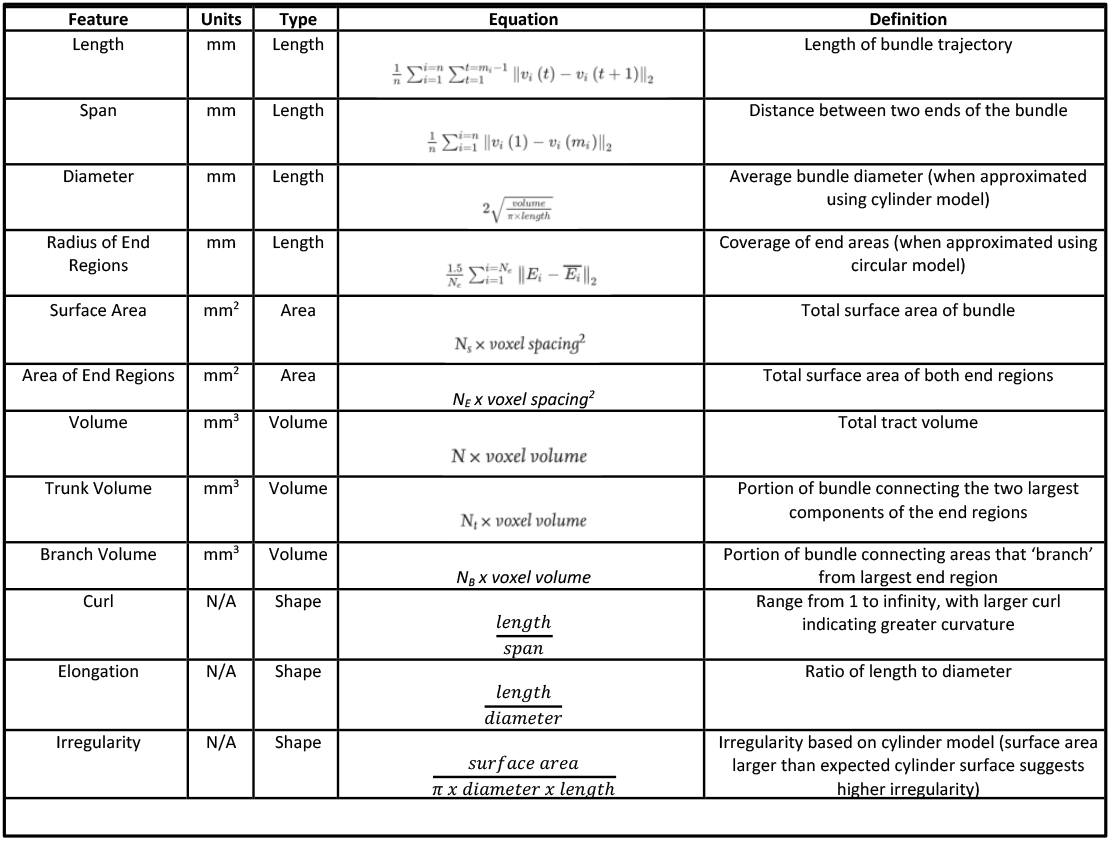
Macrostructural features, and their definitions. See (2) for complete descriptions and justifications. Following (2), a fiber bundle is a set of streamline trajectories that is represented as 3D coordinate sequences: v_i_(t) | i = 1,2,3,...n, where n is the total number of tracks, v_i_(t) is a sequence of 3D coordinates representing the trajectory of a track. t is a discrete variable from 1 to m_i_, where m_i_ is the number of the coordinates.

Quality control (QC) was performed to minimize possible false results due to acquisition issues or failure of tractography. For acquisition related QC, sessions were removed from analysis if the diffusion weighted correlation was less than 3 standard deviations away from the mean correlation (for each dataset), or if signal slice dropout occurred in >10% of slices (~3 slices). Individual bundles were removed from the analysis if the number of segmented streamlines was less than 3 standard deviations away from the mean number (for each pathway), or if the total number of streamlines was below 200 (indicating failure of tractography), and subjects were removed from analysis if >20% of pathways failed QC. We note that this stringent QC still resulted in N>2000 samples for all but 7 pathways. The total number of samples per dataset is given in supplementary data (**Supplementary Table 1**), and a list of abbreviations for all 120 (71 + 49) pathways is given in the appendix.

### Analytical Plan

To investigate the relationship between age and each WM feature, linear mixed effects modeling was performed, with each (z-normalized) feature, Y, modeled as a linear function of age, *y* = *β*_0_ + *β*_1_*Age* + *β*_2_*Sex* + *β*_3_*TICV* + *β*_3_(1 + *AGE* | *DATASET*) + *β*_4_(*SUB*), where subjects (SUB) were entered as a random effect (i.e., subject-specific random intercept), and subject sex (*Sex*) and total intracranial volume (TICV) as a fixed effects. Additionally, we modelled the association between age and outcome variable as dataset (DATASET) specific due to expected differences in MR protocols (32–36), and included a dataset specific random slope and intercept. We note that the TICV utilized was calculated from the T1-weighted image from the baseline scan, and is scaled appropriately depending on units of the feature, Y (scaled by TICV/4pi)^(1/2) for area, and scaled by (3*TICV/4pi)^(1/3 for length).

Due to multiple comparisons, all statistical tests were controlled by the false discovery rate at 0.05 to determine significance. All results are presented as the beta coefficient of estimate *‘B_1_*’, or in other words “the association of the feature ‘*y*’ with *Age”,* which (due to normalization) represents the standard deviation change in feature per year. These measures are derived for each pathway and each feature. Supplementary results additionally show the results as a percent change per year, derived from the slope normalized by the average value across the aging population (from 50-97), and multiplied by 100, which represents the percent change in feature per year.

## Results

### Total Intracranial Volume, White Matter, Gray Matter, and CSF

**Supplementary Figure 2** shows results of global changes in tissue volume. Total GM and WM tissue volumes decrease, along with increases in CSF volumes, in agreement with the literature. While GM, CSF, and global tissue volume are not the primary aims of this study, we did find significant age associations with these measures.

### What changes and where?

To summarize association with age for all features and all pathways, we show the beta coefficient associations with age for all features in matrix form in **Figure 2**, along with boxplots highlighting the percent change for all microstructure and all macrostructure features. Similar results, but shown as the percent-change-per-year from linear mixed effects models, are shown in **Supplementary Figure 3**.

**Figure 2.**
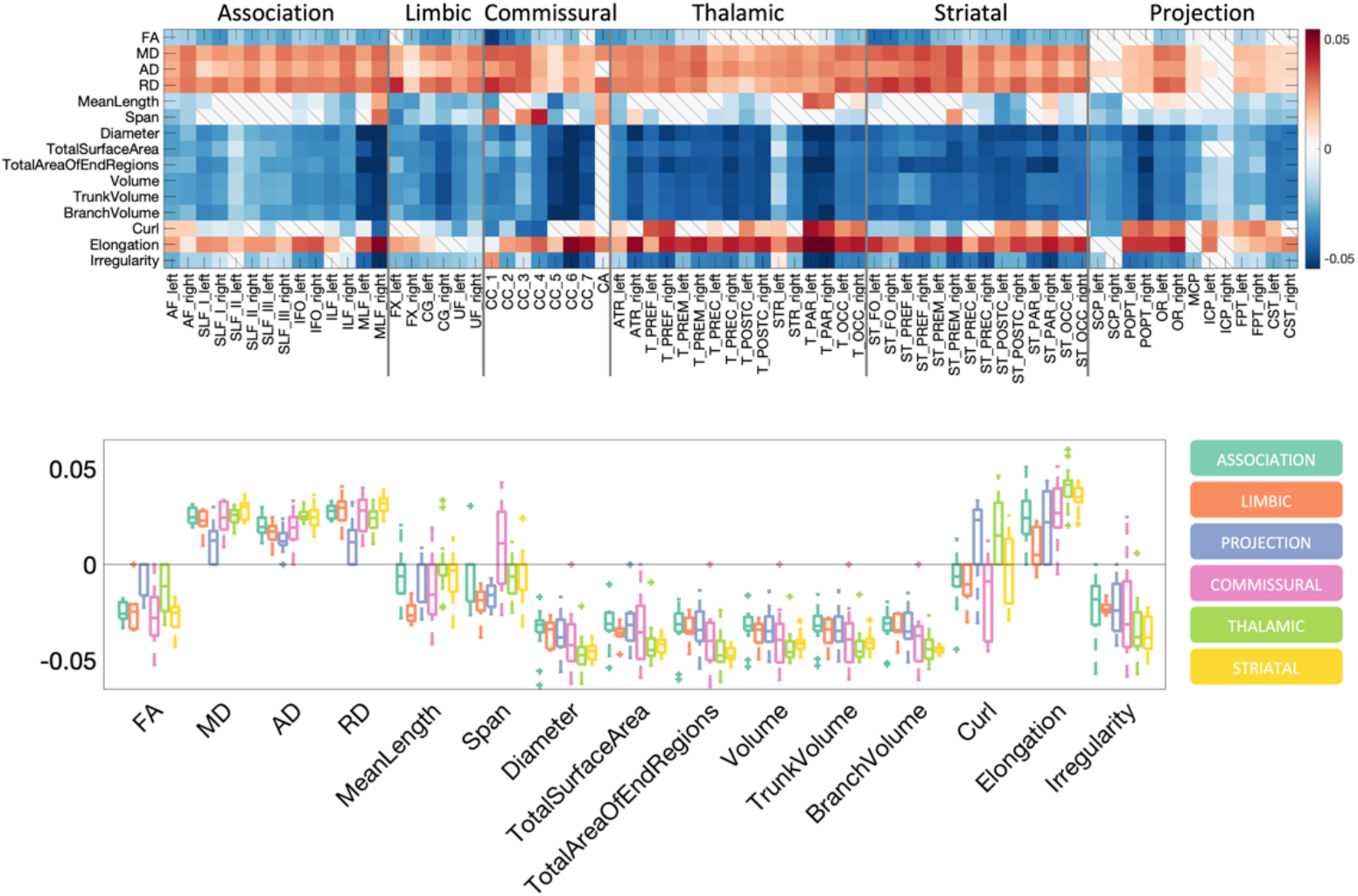
What and where changes occur during aging. The beta coefficient from linear mixed effects modeling is shown as a matrix for all features across all pathways, and also shown as boxplots for both microstructural features (left) and macrostructural features (right). Boxplots are shown separated by pathway types. Results are shown for TractSeg-derived pathways.

Most notably, microstructure measures show fairly homogenous changes across all pathways, with negative associations for FA, and again positive associations for diffusivities, with median association coefficients with age of −0.02, and approximately +0.02 to +0.03, respectively (changes of −0.2% per year, and +0.3-0.5% per year, respectively). In general, features of length, area, and volumes decrease with age, however, changes are heterogenous across pathways. Measures of volume (total volume, trunk volume, branch volume) show median associations across pathways of −0.4, −0.4, and −0.4 (changes of −0.9%, −0.9%, and −0.6% per year). Elongation show positive trends with age, while irregularity decreases with age.

Large commissural pathways (the body, splenium, and genu of the corpus callosum), as well as thalamic and striatal projections show the strongest negative trends of all features of size with age. Additionally, a number of association fibers and fasciculi, including the SLF sub-components, ILF, FAT, MLF, and PAT of both hemispheres show trends with age for all shape features, with greater changes in volumes and area of end regions than mean lengths and spans.

**Supplementary Figure 4** and **Supplementary Figure 5** show results from the ATR fiber tractography (for fit coefficients and percent-change per year, respectively), which indicate similar changes with age and in similar locations, with fit coefficients and percent-change per year of similar magnitudes. FA shows negative associations with age, diffusivities show positive associations, with microstructure measures associations similar across all pathways. Measures of volume show the greatest negative associations with age, with larger changes in the commissural and thalamic pathways.

### Visualizing change

To visualize where these changes occur, **Figure 3** shows example streamlines, separated into association, limbic, commissural, thalamic, projection, and striatal pathways, with bundles colored using the previous colormaps, and only showing bundles with statistically significant changes (**Figure 3** shows the Beta coefficients from linear mixed-effects models, **Supplementary Figure 6** shows results interpreted as percent-change per year). Notably, the changes in FA and MD are similar, with the CST changing the least (yet still statistically significant) with age, and the forceps major and anterior thalamic and striatal radiations, which occupy a majority of frontal lobe white matter space, changing the most. Other pathways show relatively homogenous change across age. Volumes and End Region Areas show similar trends, with large changes in the frontal lobe pathways, large changes in white matter of the occipital lobe, and small (but statistically significant) changes in the pathways associated with motor and pre-motor regions. The mean length decreases at a much smaller rate per year, remaining statistically significant, with visual exceptions of AC (a small commissural pathway), and projection pathways (including striatal and thalamic) to the occipital lobe. Similarly, the left and right OR show increased length with age, which would be an intuitive result of increased CSF (i.e., larger ventricles), and thus a more tortuous path from occipital lobe to thalamus. Similar results, in similar locations, are confirmed using pathways segmented using ATR, and are shown in **Supplementary Figure 7** (as Beta coefficients) and **Supplementary Figure 8** (as percent-change per year).

**Figure 3.**
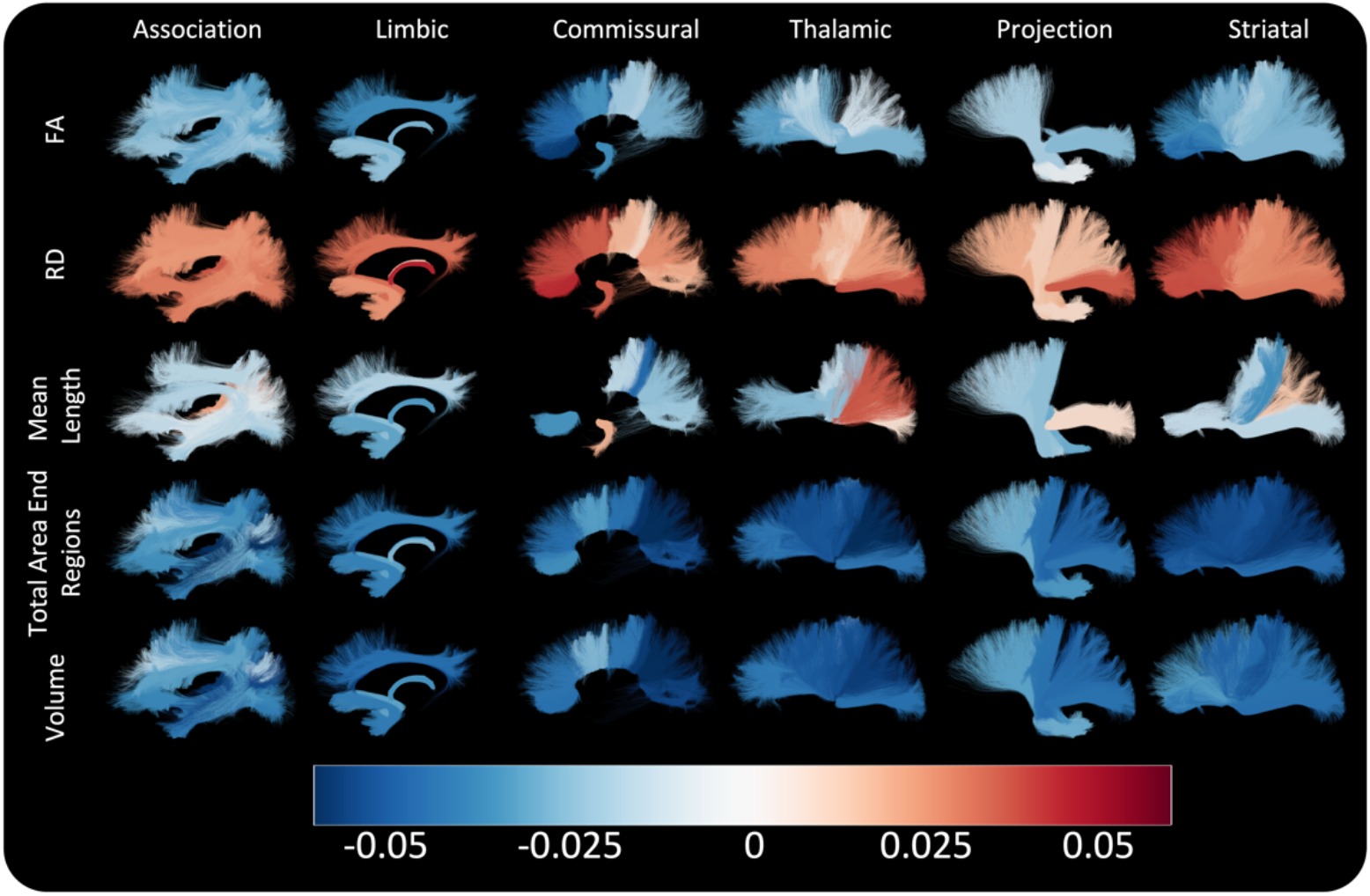
Bundle-based visualization of associations with age. Bundles that have significant associations with age are colored based on Beta-association coefficient from linear mixed-effects models, for 5 selected features. Only those with statistically significant change with age are displayed. Results are shown for TractSeg-derived pathways.

### Pathways of interest

To provide even more insight into the microstructural and macrostructural associations shown in this study, we have provided illustrations for a projection tract (i.e., anterior thalamic radiation, **Figure 4**) and commissural tract (i.e., forceps minor, **Figure 5**). For the anterior thalamic radiation (3D illustration in **Figure 4A**), we found significant age-related decline in all four microstructural measures (**Figure 4B**), in which there was a positive age-related association with MD (p=3E-5), RD (p=5E-5), and AD (p=6E-4), and a negative association with FA (p=3E-4). There were also several significant associations with macrostructural features for this tract. **Figure 4C** illustrates 4 of these associations, including volume (p=1E-6), branch volume (p=3E-3), surface area (p=1E-7), and area of end regions (p=0.02). **Figure 5** illustrates the associations for the forceps minor tract, again demonstrating significant positive age-related associations with diffusivities, negative age-related associations with FA, and negative age-related associations with volume, surface area, and area of end regions.

**Figure 4.**
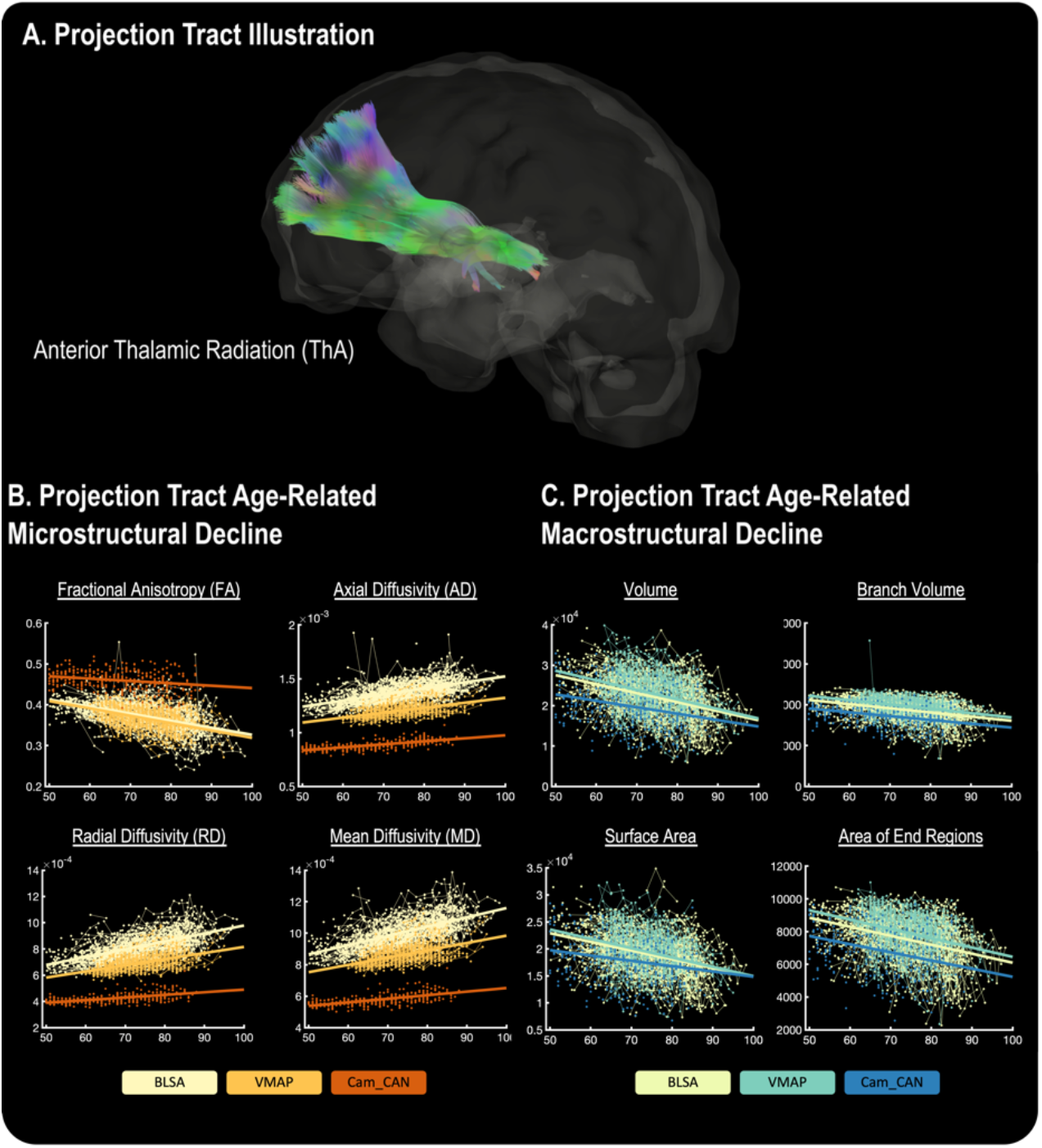
Example microstructural and macrostructural associations for a projection white matter tract. A 3D illustration of the anterior thalamic radiation (ThA) is shown (A), as it exhibited significant microstructural (B) and microstructural decline (C). For each microstructural and macrostructural plot, colored datapoints and lines represent individual cohorts.

**Figure 5.**
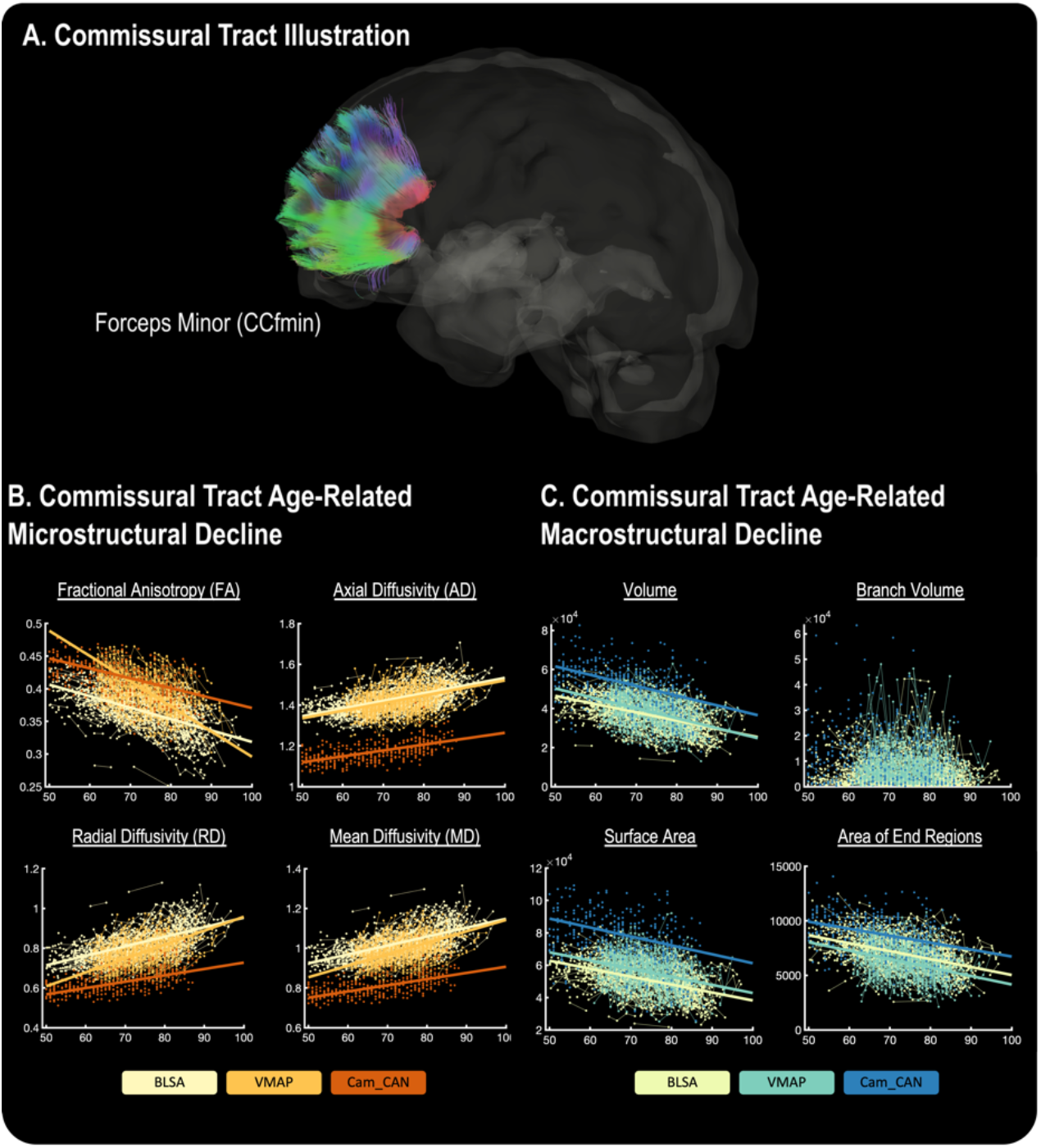
Example microstructural and macrostructural associations for a commissural white matter tract. A 3D illustration of the forceps minor (CCfmin) is shown (A), as it exhibited significant microstructural (B) and microstructural decline (C). For each microstructural and macrostructural plot, colored datapoints and lines represent individual cohorts.

## Discussion

Using a large, cross sectional and longitudinal dataset, we analyze microstructural features and, for the first time, shape-based features, of WM pathways across age. While prior studies have evaluated how WM microstructure changes with age, few studies have determined if these patterns differ across different types of WM tracts. The novelty of this study is that we evaluated the heterogeneous age-related decline in microstructural and macrostructural features in four types of WM tracts: association, limbic, projection, and commissural. We found that while microstructural features were globally sensitive to age-related decline, these measures were largely homogeneous in their decline across the association, limbic, projection, and commissural fibers. In contrast, we found that macrostructural features were non-uniform in their trends in age-related decline. Specifically, we found that, overall, the projection and commissural fibers demonstrated more age-related decline than the association and limbic fibers. Thus, macrostructural features may be more specific in identifying age-related WM decline, and could a more sensitive marker for neurodegenerative disorders compared to microstructural features.

### Age-Related Microstructural Decline

Trends seen in diffusion microstructure indices mirror that from existing literature, which we have confirmed generalize to larger datasets, and across datasets with different scanners, vendors, and acquisitions. Diffusivities increase with age, with the largest change shown for radial, and mean diffusivities, and to a lesser extent, axial diffusivities. Consequently, this leads to a decrease in fractional anisotropy. This has traditionally been attributed to myelin loss and/or decreased axonal volume fractions and densities (17, 21, 37, 38), with supplemental evidence provided through advanced multicompartment modeling (18, 21). However, care must be taken when interpreting these indices as highly specific markers of tissue microstructure, as diffusion (and DTI in particular) is sensitive to a number of potential biophysical changes (39).

As expected, the datasets showed large effects on quantified measures (36, 40) due to differences in acquisition conditions (33, 34, 41), although the same trends were seen across datasets, with only small differences in associations with age. Combination of datasets in analysis requires either accounting for these effects in modeling (as performed here) or harmonizing data across scanners and sites, which is an active area of interest (36, 42, 43). Harmonization studies can utilize these well-characterized effects of age as validation of techniques and algorithms.

### Age-Related Macrostructural Decline

While tractography has been used to study the human brain in aging, it is often used to simply extract pathway-specific indices of microstructure (or quantitative) measures. Here, we study shape-based features of tractography-defined bundles, quantifying basic features (e.g., length, diameter, volume) and more comprehensive features (e.g., curl, irregularity, elongation). We find that, indeed, the shape of white matter features changes with age. Notably, basic macrostructural features like volume and total surface areas exhibit age-related decline, in agreement with the observed trend of a decrease in total white matter volume. Further, more comprehensive measures, such as irregularity and elongation, age-related decline. In our subsequent analysis to determine if there was heterogeneous age-related decline between the association, limbic, association, and commissural tracts, we found widespread significant differences (see ***Figure 2***). For example, age-related changes in elongation (length ÷ diameter) were relatively low in the association and limbic tracts but higher in the projection and commissural tracts. Furthermore, age-related changes were similar for irregularity [surface area ÷ (π x diameter x length)], in which age-related decline was lower in the projection and commissural tracts compared to the association and limbic tracts.

Our findings therefore indicate that specific white matter features can be used to identify age-related decline, and these features can also be incorporated into clinical populations to identify abnormal aging patterns. Future work should investigate different trends in disease cohorts, where this analysis facilitates asking “what changes?” and “where?”. This also results in the creation of a large feature space (10’s of pathways x 10’s of features) for each subject, which may facilitate machine learning, deep learning, and dimensionality reduction techniques to identify abnormalities in an individual subject or cohort. Similar analysis may also be used in an unsupervised fashion – rather than utilizing predefined bundles, a connectome-style approach can be used to extract every fiber bundle in a large connectome matrix followed by subsequent feature-based analysis of every edge in the connectome.

### Global and local changes

In general, all pathways show consistent changes in tissue microstructure with age, indicating a largely *global* change in microstructure. In contrast, shape features of pathways show very different effect sizes and relative changes per year across the brain, which indicates *local* changes, and pathway-specific differences image. Overall, this suggests that microstructure features of pathways change together, and at relatively the same rates, whereas macrostructural features do not and indicate location-specific indices of change, whereby projection and commissural fibers exhibit more significant age-related decline. Thus, pathway features might be a more sensitive biomarker for differences due to disease or disorders.

### Limitations

This study has several limitations. While we utilized large samples sizes and showed generalizability to very different aging datasets, results were tested on just one bundle segmentation algorithm. Additionally, many pathways were investigated, significantly more than is typical for many studies on aging, and many of these pathways are smaller association pathways that may be harder or more variable to track. Nevertheless, the large sample size facilitated statistical analysis and findings with small effect sizes. The use of different datasets with different acquisitions is known to result in very different quantitative indices, and in the current study, very different tractography results. However, we consider this an advantage to the current study where results generalized across all data, and effect of dataset was included in modeling.

Future studies should investigate and characterize shape changes across the lifespan. This may be particularly relevant in childhood where large changes in brain structure and microstructure are expected. Second, the combination of shape and microstructure features in disease should be investigated. There is a significant body of research on DTI changes in disease, and it is intuitive that the shape, location, and geometry of pathways may also experience significant alterations in such states. Finally, the relationship between GM regions and WM structure should be investigated. The full feature space of GM volume, thickness, and surface area, in combination with WM macrostructure and microstructure features, will facilitate a complete description of changes in the brain during aging.

## Conclusion

We provide a comprehensive characterization of WM changes in aging. Using large cross-sectional and longitudinal diffusion datasets, we have shown that both microstructural and macrostructural geometrical features of the human brain change during normal healthy aging. Microstructural indices of anisotropy and diffusivity show the largest effects with age, with global trends apparent across all pathways. Macrostructural features of volume, surfaces areas, and lengths also change with age, with trends that are not uniform across all pathways. Thus, tract-specific changes in geometry occur in normal aging. Results from this study may be useful in understanding biophysical and structural changes occurring during normal aging and will facilitate comparisons in a variety of diseases or abnormal conditions.

## Statements and Declarations

### Author Contributions

All authors contributed to the study conception and design. Data collection was performed by the Baltimore Longitudinal Study of Aging at the National Institutes of Aging, and the Vanderbilt Memory & Aging Project (VMAP). All authors commented on previous versions of the manuscript. All authors read and approved the final manuscript.

### Data Availability

The summary microstructure and macrostructure features, for all pathways and subjects, along with demographic information, will be made available upon acceptance of the manuscript.

### Ethic Approval

All human datasets from Vanderbilt University were acquired after informed consent under supervision of the appropriate Institutional Review Board. All additional datasets are freely available and unrestricted for noncommercial research purposes. This study accessed only deidentified patient information.

### Consent to participate

Informed consent was obtained from all individual participants included in the study.

## Appendix

The bundles resulting from each segmentation pipeline are given as a list below, with acronyms used in the text.

### TractSeg

Arcuate fascicle left (AF_L); Arcuate fascicle right (AF_R); Anterior Thalamic Radiation left (ATR_L); Thalamic Radiation right; (ATR_R); Commissure Anterior (CA); Rostrum (CC_1; Genu (CC_2); Rostral body (Premotor) (CC_3); Anterior midbody (Primary Motor) (CC_4); Posterior midbody (Primary Somatosensory) (CC_5); Isthmus (CC_6); Splenium (CC_7); Corpus Callosum –all (CC); Cingulum left (CG_L); Cingulum right (CG_R); Corticospinal tract left (CST_L); Corticospinal tract right (CST_R); Fronto-pontine tract left (FPT_L); Fronto-pontine tract right (FPT_R); Fornix left (FX_L); Fornix right (FX_R); Inferior cerebellar peduncle left (ICP_L); Inferior cerebellar peduncle right (ICP_R); In-ferior occipito-frontal fascicle left (IFO_L); Inferior occipito-frontal fascicle right (IFO_R); Inferior longitudinal fascicle left (ILF_L); Inferior longitudinal fascicle right (ILF_R); Middle cerebellar peduncle (MCP); Middle longitudinal fascicle left (MLF_L); Middle longitudinal fascicle right (MLF_R); Optic radiation left (OR_L); Optic radiation right (OR_R); Parieto-occipital pontine left (POPT_L); Parieto-occipital pontine right (POPT_R); Superior cerebellar peduncle left (SCP_L); Superior cerebellar peduncle right (SCP_R); Superior longitudinal fascicle III left SLF_III_L); Superior longitudinal fascicle III right (SLF_III_R); Superior longitudinal fascicle II left (SLF_II_L); Superior longitudinal fascicle II right (SLF_II_R); Superior longitudinal fascicle I left (SLF_I_L); Superior longitudinal fascicle I right (SLF_I_R); Striato-fronto-orbital left (ST_FO_L); Striato-fronto-orbital right (ST_FO_R); Striato-occipital left (ST_OCC_L); Striato-occipital right (ST_OCC_R); Striato-parietal left (ST_PAR_L); Striato-parietal right (ST_PAR_R); Striato-postcentral left (ST_POSTC_L); Striato-postcentral right (ST_POSTC_R); Striato-precentral left (ST_PREC_L); Striato-precentral right (ST_PREC_R); Striato-prefrontal left (ST_PREF_L); Striato-prefrontal right (ST_PREF_R); Striato-premotor left (ST_PREM_L); Striato-premotor right (ST_PREM_R); Thalamo-occipital left (T_OCC_L); Thalamo-occipital right (T_OCC_R); Thalamo-parietal left (T_PAR_L); Thalamo-parietal right (T_PAR_R); Thalamo-postcentral left (T_POSTC_L); Thalamo-postcentral right (T_POSTC_R); Thalamo-precentral left (T_PREC_L); Thalamo-precentral right (T_PREC_R); Thalamo-prefrontal left (T_PREF_L); Thalamo-prefrontal right (T_PREF_R); Thalamo-premotor left (T_PREM_L); Thalamo-premotor right (T_PREM_R); Uncinate fascicle left (UF_L); Uncinate fascicle right (UF_R).

### ATR

Arcuate_Fasciculus_L (AF_L); Arcuate Fasciculus R (AF_R); Cortico Spinal Tract L (CST_L); Cortico Spinal Tract R (CST_R); Cortico Striatal Pathway L (CS_L); Cortico Striatal Pathway R (CS_R); Corticobulbar Tract L (CBT_L); Corticobulbar Tract R (CBT_R); Corticopontine Tract L (CPT_L); Corticopontine Tract R (CPT_R); Corticothalamic Pathway L (CTP_L); Corticothalamic Pathway R (CTP_R); Inferior Cerebellar Peduncle L (ICP_L); Inferior Cerebellar Peduncle R (ICP_R); Inferior Fronto Occipital Fasciculus L (IFOF_L); Inferior Fronto Occipital Fasciculus R (IFOF_R); Inferior Longitudinal Fasciculus L (ILF_L); Inferior Longitudinal Fasciculus R (ILF_R); Optic Radiation L (OR_L); Optic Radiation R (OR_R); Middle Longitudinal Fasciculus L (MdLF_L); Middle Longitudinal Fasciculus R (MdLF_R); Uncinate Fasciculus L (UF_L); Uncinate Fasciculus R (UF_R).

## Supplementary Material

**Supplementary Figure 1.**
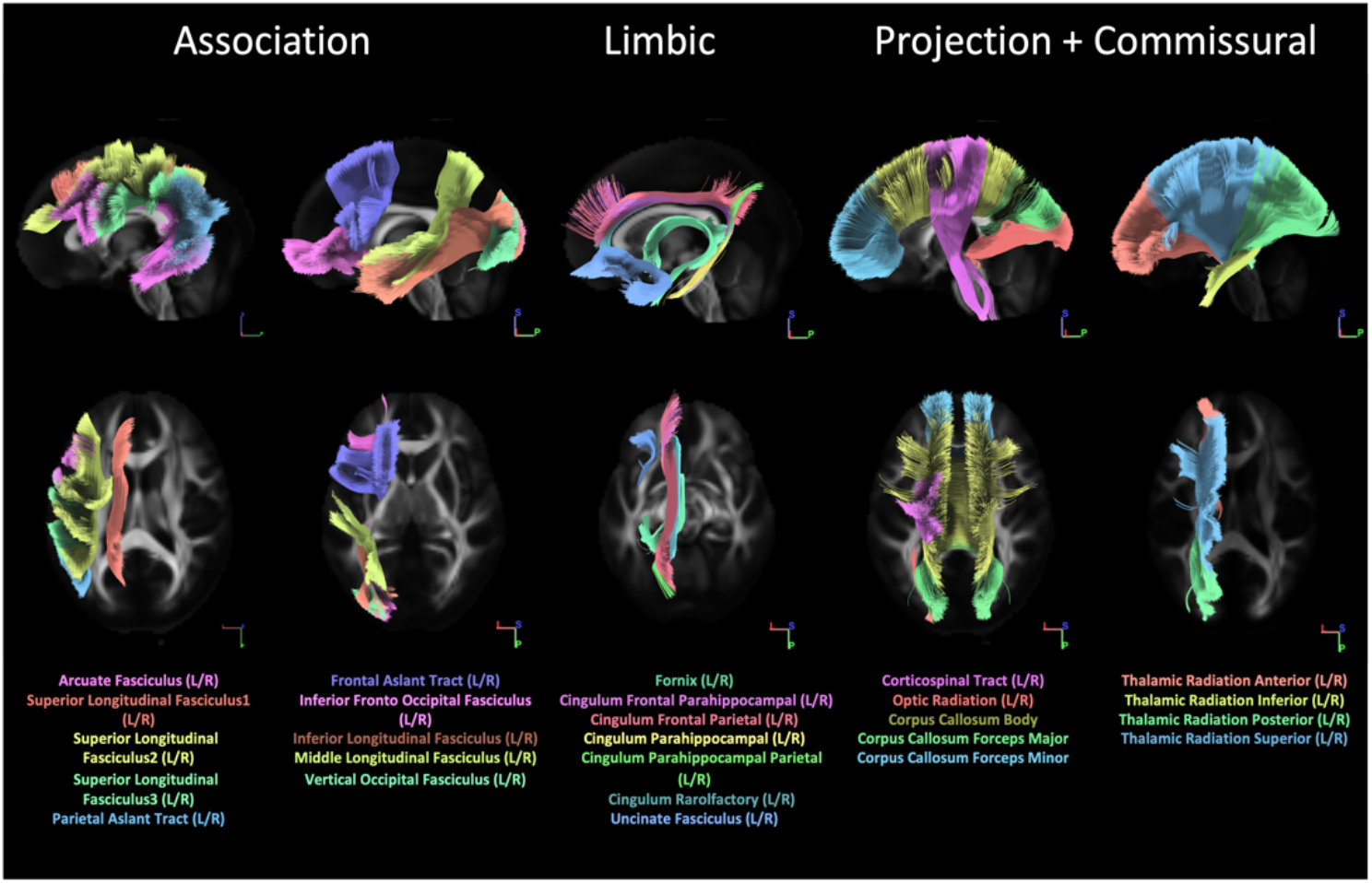
We investigated microstructure and macrostructure features of 49 pathways virtually segmented using DSI Studio, visualized and organized into association, limbic, projection, and commissural fibers.

**Supplementary Table 1.** Abbreviations and sample size for studied pathways.

| PATHWAY | ABBREVIATION | BLSA | CAMCAN | VMAP | TOTAL |
| --- | --- | --- | --- | --- | --- |
| Arcuate Fasciculus L | AF_L | 1312 | 839 | 354 | 2505 |
| Arcuate Fasciculus R | AF_R | 1184 | 743 | 301 | 2228 |
| Cingulum Frontal Parahippocampal L | CgFPH_L | 1036 | 728 | 329 | 2093 |
| Cingulum Frontal Parahippocampal R | CgFPH_R | 1077 | 734 | 340 | 2151 |
| Cingulum Frontal Parietal L | CgFP_L | 1310 | 842 | 355 | 2507 |
| Cingulum Frontal Parietal R | CgFP_R | 1319 | 843 | 354 | 2516 |
| Cingulum Parahippocampal L | CgP_L | 1313 | 841 | 353 | 2507 |
| Cingulum Parahippocampal R | CgP_R | 1318 | 839 | 352 | 2509 |
| Cingulum Parahippocampal Parietal L | CgPP_L | 850 | 643 | 240 | 1733 |
| Cingulum Parahippocampal Parietal R | CgPP_R | 1112 | 771 | 319 | 2202 |
| Cingulum Rarolfactory L | CgR_L | 431 | 198 | 200 | 829 |
| Cingulum Rarolfactory R | CgR_R | 1230 | 757 | 352 | 2339 |
| Corpus Callosum Body | CCbody | 1266 | 775 | 335 | 2376 |
| Corpus Callosum Forceps Major | CCfmaj | 1321 | 836 | 353 | 2510 |
| Corpus Callosum Forceps Minor | CCfmin | 1322 | 845 | 351 | 2518 |
| Corticospinal Tract L | CST_L | 1275 | 843 | 356 | 2474 |
| Corticospinal Tract R | CST_R | 1259 | 845 | 351 | 2455 |
| Fornix L | Fx_L | 859 | 763 | 274 | 1896 |
| Fornix R | Fx_R | 629 | 642 | 258 | 1529 |
| Frontal Aslant Tract L | FAT_L | 1303 | 835 | 355 | 2493 |
| Frontal Aslant Tract R | FAT_R | 1311 | 843 | 356 | 2510 |
| Inferior Frontal Occipital Fasciculus L | IFOF_L | 1298 | 822 | 355 | 2475 |
| Inferior Frontal Occipital Fasciculus R | IFOFs_R | 1316 | 827 | 354 | 2497 |
| Inferior Longitudinal Fasciculus L | ILF_L | 1309 | 842 | 355 | 2506 |
| Inferior Longitudinal Fasciculus R | ILF_R | 1321 | 834 | 355 | 2510 |
| Middle Longitudinal Fasciculus L | MLF_L | 1211 | 738 | 335 | 2284 |
| Middle Longitudinal Fasciculus R | MLF_R | 1249 | 722 | 328 | 2299 |
| Optic Radiation L | OR_L | 1140 | 783 | 354 | 2277 |
| Optic Radiation R | OR_R | 930 | 592 | 355 | 1877 |
| Parietal Aslant Tract L | PAT_L | 1319 | 840 | 354 | 2513 |
| Parietal Aslant Tract R | PAT_R | 1322 | 843 | 355 | 2520 |
| Superior Longitudinal Fasciculus1 L | SLF1_L | 1317 | 841 | 355 | 2513 |
| Superior Longitudinal Fasciculus1 R | SLF1_R | 1313 | 833 | 351 | 2497 |
| Superior Longitudinal Fasciculus2 L | SLF2_L | 1319 | 844 | 355 | 2518 |
| Superior Longitudinal Fasciculus2 R | SLF2_R | 1243 | 799 | 345 | 2387 |
| Superior Longitudinal Fasciculus3 L | SLF3_L | 1319 | 841 | 354 | 2514 |
| Superior Longitudinal Fasciculus3 R | SLF3_R | 1310 | 843 | 355 | 2508 |
| Thalamic Radiation Anterior L | ThA_L | 964 | 759 | 346 | 2069 |
| Thalamic Radiation Anterior R | ThA_R | 1069 | 764 | 352 | 2185 |
| Thalamic Radiation Inferior L | ThI_L | 1303 | 831 | 353 | 2487 |
| Thalamic Radiation Inferior R | ThI_R | 1155 | 788 | 353 | 2296 |
| Thalamic Radiation Posterior L | ThP_L | 1184 | 788 | 353 | 2325 |
| Thalamic Radiation Posterior R | ThP_R | 1088 | 592 | 344 | 2024 |
| Thalamic Radiation Superior L | ThS_L | 633 | 397 | 348 | 1378 |
| Thalamic Radiation Superior R | ThS_R | 453 | 183 | 353 | 989 |
| Uncinate Fasciculus L | UF_L | 1320 | 839 | 352 | 2511 |
| Uncinate Fasciculus R | UF_R | 1250 | 768 | 352 | 2370 |
| Vertical Occipital Fasciculus L | VOF_L | 1321 | 839 | 353 | 2513 |
| Vertical Occipital Fasciculus R | VOF_R | 1316 | 833 | 356 | 2505 |

**Supplementary Figure 2.**
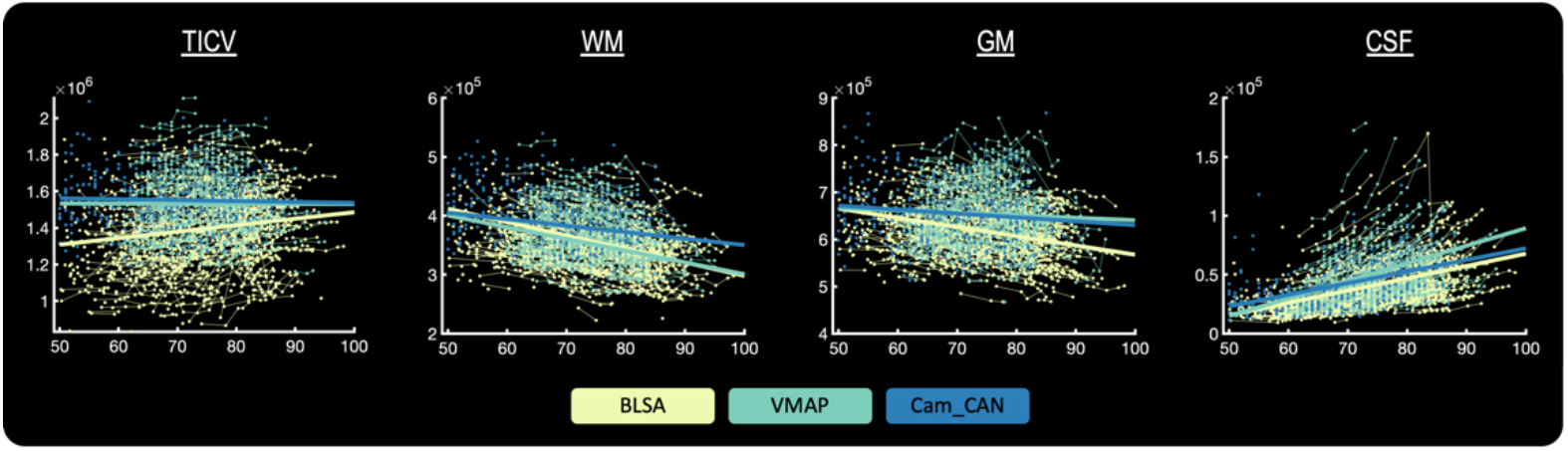
Global changes in tissue volume occur in aging, including decreases in GM volume, and WM volume, and increase in CSF volume. Colors of datapoints and lines represent 3 different datasets utilized.

**Supplementary Figure 3.**
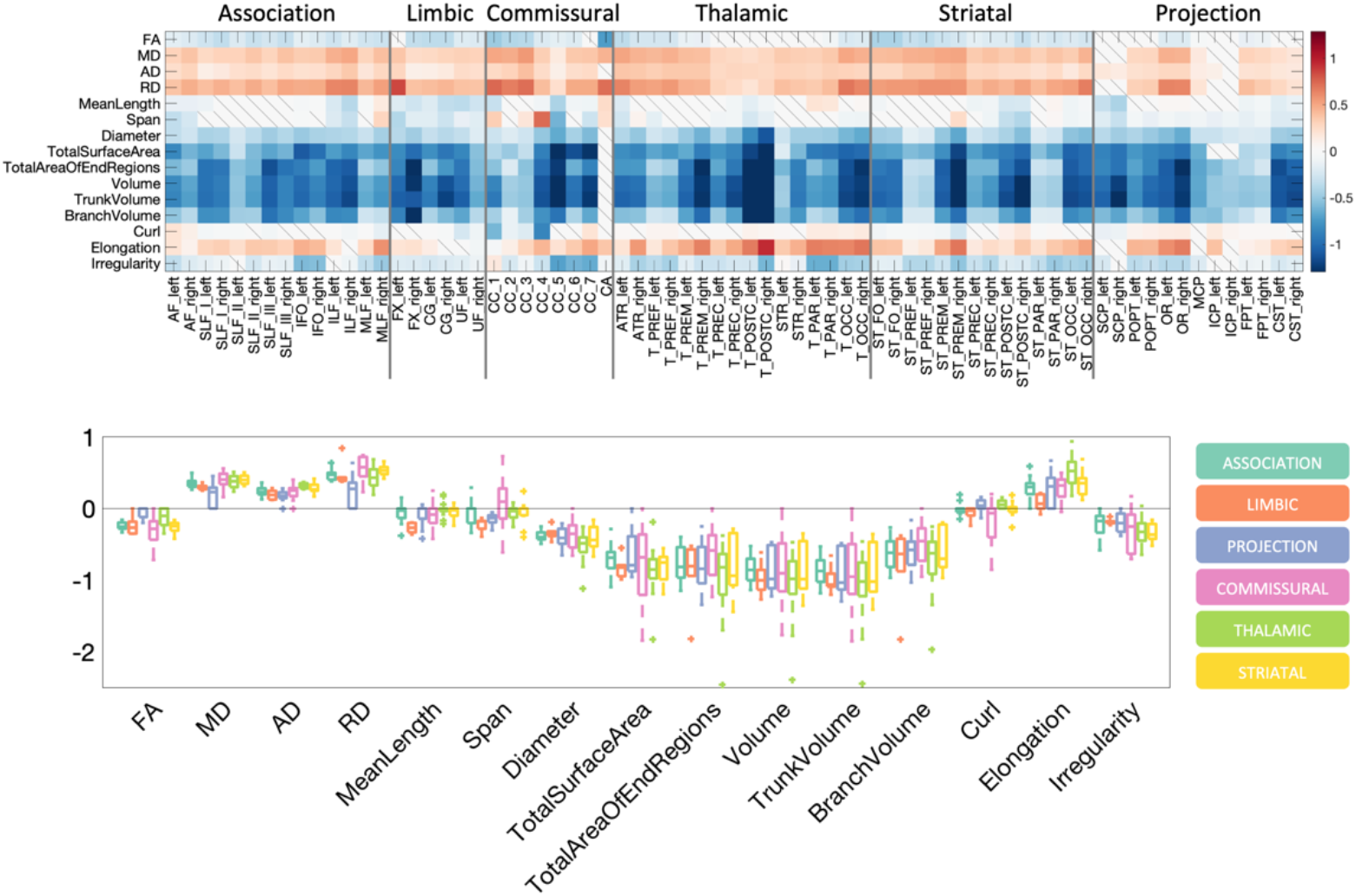
What and where changes occur during aging. The percent-change per year is shown as a matrix for all features across all pathways, and also shown as boxplots for both microstructural features (left) and macrostructural features (right). Boxplots are shown separated by pathway types. Results are shown for TractSeg-derived Pathways.

**Supplementary Figure 4.**
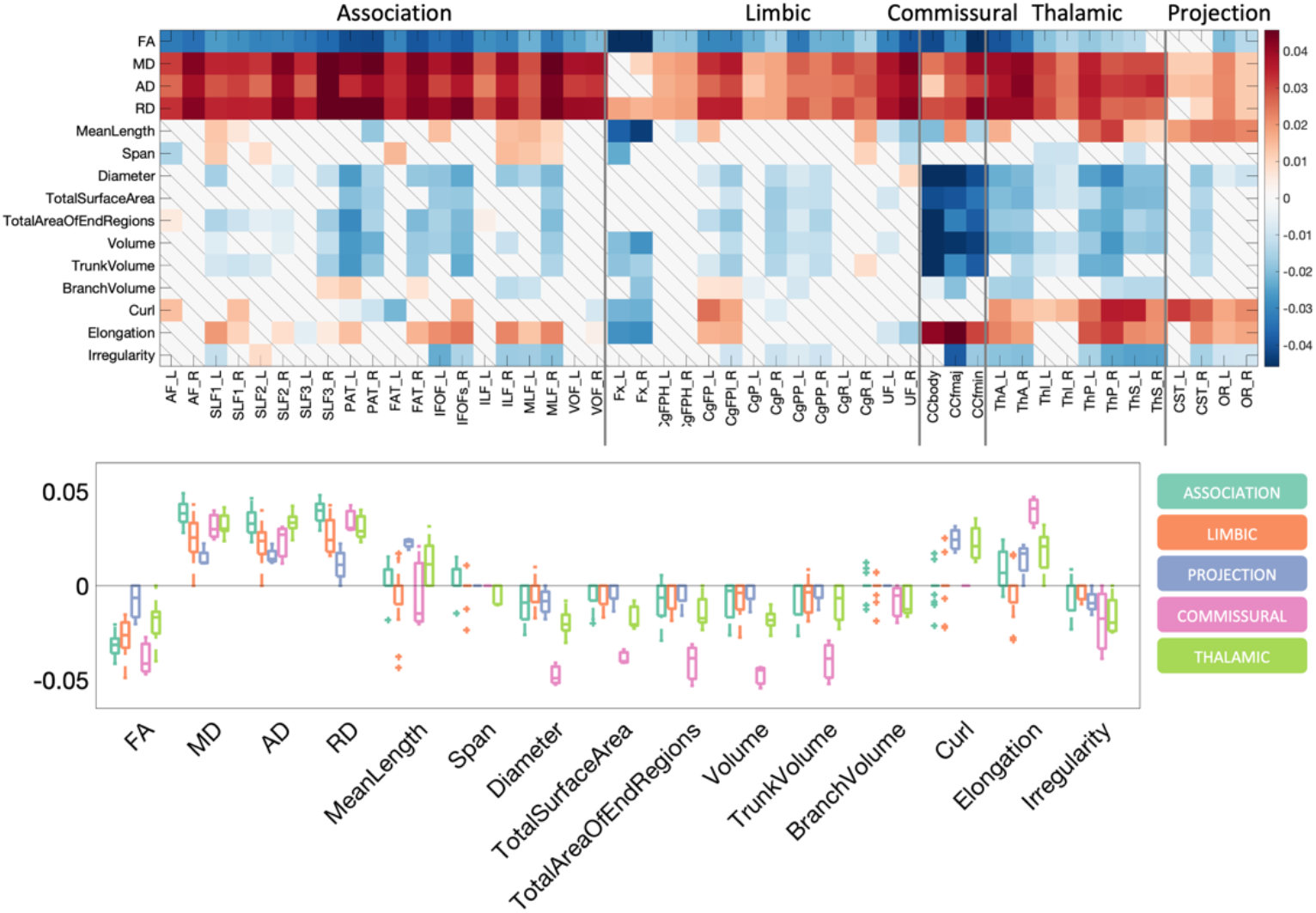
What and where changes occur during aging. The beta coefficient from linear mixed effects modeling is shown as a matrix for all features across all pathways, and also shown as boxplots for both microstructural features (left) and macrostructural features (right). Boxplots are shown separated by pathway types. Results are shown for ATR-derived pathways.

**Supplementary Figure 5.**
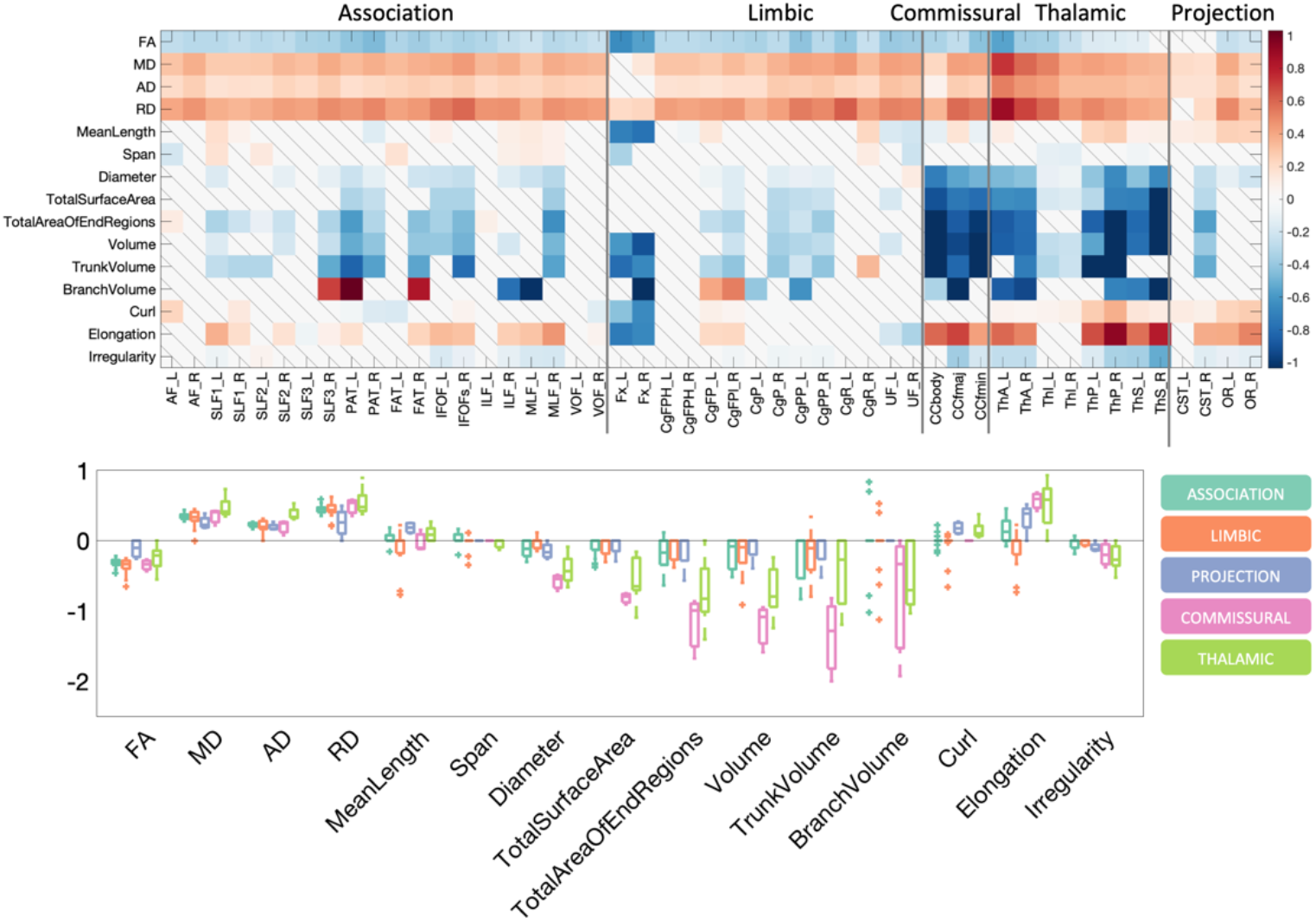
What and where changes occur during aging. The percent-change per year is shown as a matrix for all features across all pathways, and also shown as boxplots for both microstructural features (left) and macrostructural features (right). Boxplots are shown separated by pathway types. Results are shown for ATR-derived Pathways.

**Supplementary Figure 6.**
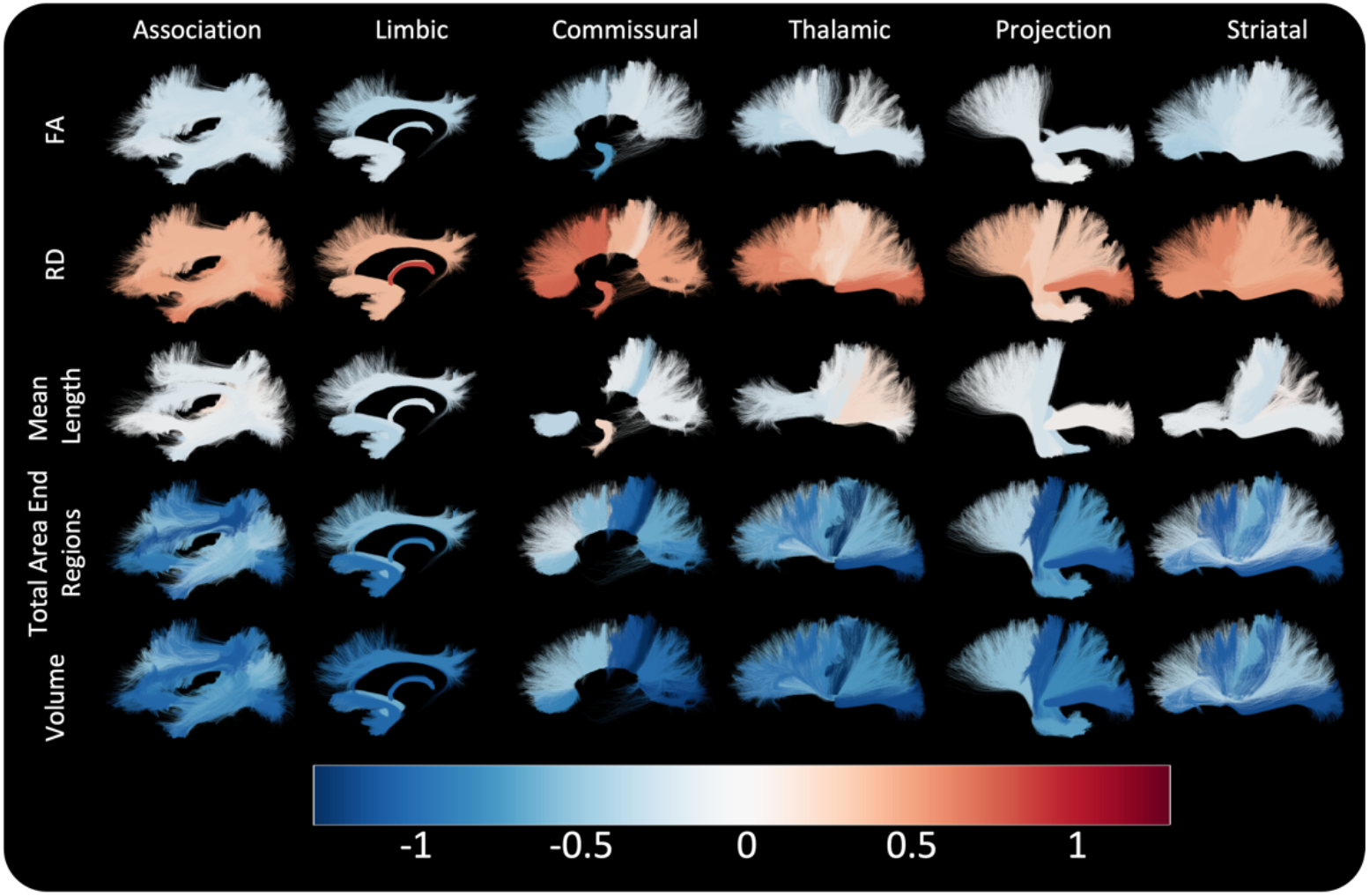
Bundle-based visualization of changes during aging. Bundles that have significant associations with age are colored based on percent-change per year, for 5 selected features. Only those with statistically significant change with age are displayed. Results are shown for TractSeg-derived pathways.

**Supplementary Figure 7.**
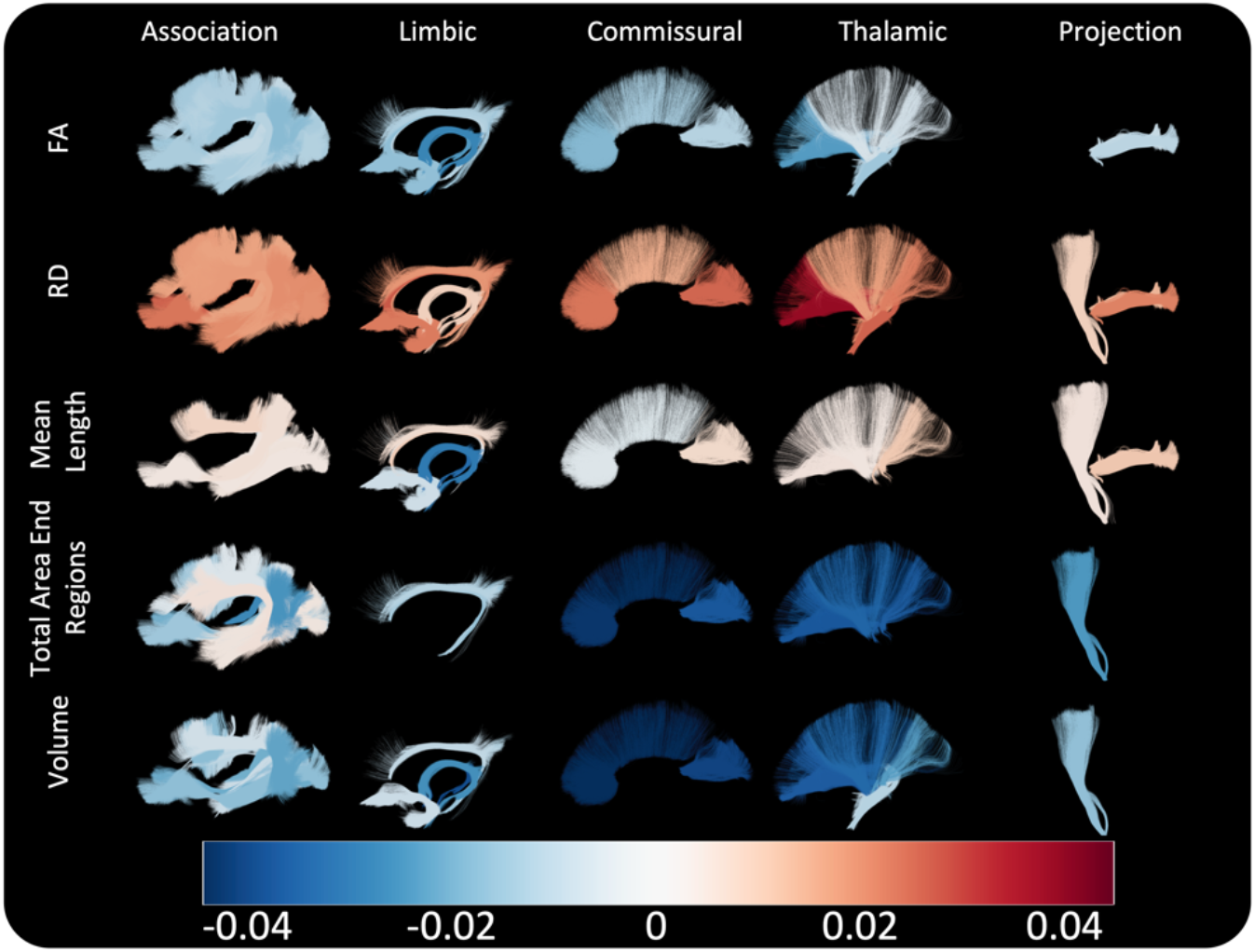
Bundle-based visualization of associations with age. Bundles that have significant associations with age are colored based on Beta-association coefficient from linear mixed-effects models, for 5 selected features. Only those with statistically significant change with age are displayed. Results are shown for ATR-derived pathways.

**Supplementary Figure 8.**
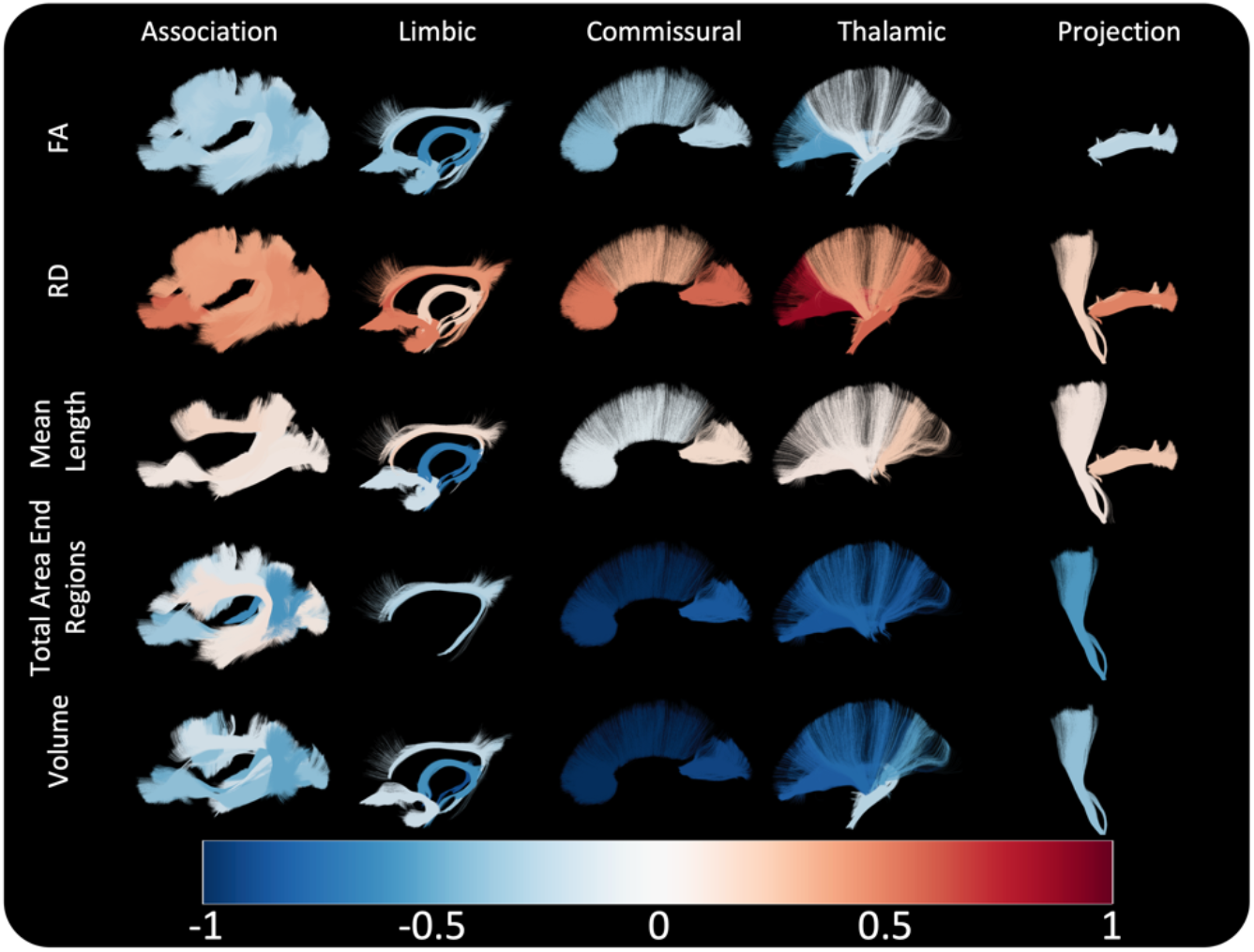
Bundle-based visualization of changes during aging. Bundles that have significant associations with age are colored based on percent-change per year, for 5 selected features. Only those with statistically significant change with age are displayed. Results are shown for ATR-derived pathways.

## References

1. J. Wasserthal, P. Neher, K. H. Maier-Hein, TractSeg - Fast and accurate white matter tract segmentation. Neuroimage 183, 239–253 (2018).

2. F.-C. Yeh, Shape Analysis of the Human Association Pathways. bioRxiv 10.1101/2020.04.19.049544, 2020.2004.2019.049544 (2020).

3. S. Ramanoel et al., Gray Matter Volume and Cognitive Performance During Normal Aging. A Voxel-Based Morphometry Study. Front Aging Neurosci 10, 235 (2018).

4. D. Terribilli et al., Age-related gray matter volume changes in the brain during non-elderly adulthood. Neurobiol Aging 32, 354–368 (2011).

5. K. L. Bergfield et al., Age-related networks of regional covariance in MRI gray matter: reproducible multivariate patterns in healthy aging. Neuroimage 49, 1750–1759 (2010).

6. Y. Taki et al., Correlations among brain gray matter volumes, age, gender, and hemisphere in healthy individuals. PLoS One 6, e22734 (2011).

7. A. Giorgio et al., Age-related changes in grey and white matter structure throughout adulthood. Neuroimage 51, 943–951 (2010).

8. N. Zuo et al., Gray Matter-Based Age Prediction Characterizes Different Regional Patterns. Neurosci Bull 37, 94–98 (2021).

9. A. Pfefferbaum et al., Brain gray and white matter volume loss accelerates with aging in chronic alcoholics: a quantitative MRI study. Alcohol Clin Exp Res 16, 1078–1089 (1992).

10. C. L. Kimmel et al., Age-related parieto-occipital and other gray matter changes in borderline personality disorder: A meta-analysis of cortical and subcortical structures. Psychiatry Res Neuroimaging 251, 15–25 (2016).

11. J. Wang et al., Gray Matter Age Prediction as a Biomarker for Risk of Dementia. Proc Natl Acad Sci U S A 116, 21213–21218 (2019).

12. L. Jorge et al., Investigating the Spatial Associations Between Amyloid-beta Deposition, Grey Matter Volume, and Neuroinflammation in Alzheimer’s Disease. J Alzheimers Dis 80, 113–132 (2021).

13. Y. Guo et al., Grey-matter volume as a potential feature for the classification of Alzheimer’s disease and mild cognitive impairment: an exploratory study. Neurosci Bull 30, 477–489 (2014).

14. O. Abe et al., Aging in the CNS: comparison of gray/white matter volume and diffusion tensor data. Neurobiol Aging 29, 102–116 (2008).

15. A. B. Storsve, A. M. Fjell, A. Yendiki, K. B. Walhovd, Longitudinal Changes in White Matter Tract Integrity across the Adult Lifespan and Its Relation to Cortical Thinning. PLoS One 11, e0156770 (2016).

16. Q. J. Yap et al., Tracking cerebral white matter changes across the lifespan: insights from diffusion tensor imaging studies. J Neural Transm (Vienna) 120, 1369–1395 (2013).

17. C. Lebel et al., Diffusion tensor imaging of white matter tract evolution over the lifespan. Neuroimage 60, 340–352 (2012).

18. D. Beck et al., White matter microstructure across the adult lifespan: A mixed longitudinal and crosssectional study using advanced diffusion models and brain-age prediction. Neuroimage 224, 117441 (2021).

19. N. Toschi, R. A. Gisbert, L. Passamonti, S. Canals, S. De Santis, Multishell diffusion imaging reveals sex-specific trajectories of early white matter degeneration in normal aging. Neurobiol Aging 86, 191–200 (2020).

20. Y. S. Chang et al., White Matter Changes of Neurite Density and Fiber Orientation Dispersion during Human Brain Maturation. PLoS One 10, e0123656 (2015).

21. S. R. Cox et al., Ageing and brain white matter structure in 3,513 UK Biobank participants. Nat Commun 7, 13629 (2016).

22. W. Y. Isaac Tseng et al., Microstructural differences in white matter tracts across middle to late adulthood: a diffusion MRI study on 7167 UK Biobank participants. Neurobiol Aging 98, 160–172 (2021).

23. O. A. Williams et al., Vascular burden and APOE epsilon4 are associated with white matter microstructural decline in cognitively normal older adults. Neuroimage 188, 572–583 (2019).

24. A. L. Jefferson et al., The Vanderbilt Memory & Aging Project: Study Design and Baseline Cohort Overview. J Alzheimers Dis 52, 539–559 (2016).

25. J. R. Taylor et al., The Cambridge Centre for Ageing and Neuroscience (Cam-CAN) data repository: Structural and functional MRI, MEG, and cognitive data from a cross-sectional adult lifespan sample. Neuroimage 144, 262–269 (2017).

26. F. C. Yeh et al., Population-averaged atlas of the macroscale human structural connectome and its network topology. Neuroimage 178, 57–68 (2018).

27. K. G. Schilling et al., Tractography dissection variability: What happens when 42 groups dissect 14 white matter bundles on the same dataset? Neuroimage 243, 118502 (2021).

28. J. D. Tournier et al., MRtrix3: A fast, flexible and open software framework for medical image processing and visualisation. Neuroimage 202, 116137 (2019).

29. F. C. Yeh, V. J. Wedeen, W. Y. Tseng, Generalized q-sampling imaging. IEEE Trans Med Imaging 29, 1626–1635 (2010).

30. F. C. Yeh, T. D. Verstynen, Y. Wang, J. C. Fernandez-Miranda, W. Y. Tseng, Deterministic diffusion fiber tracking improved by quantitative anisotropy. PLoS One 8, e80713 (2013).

31. F. C. Yeh et al., Automatic Removal of False Connections in Diffusion MRI Tractography Using Topology-Informed Pruning (TIP). Neurotherapeutics 16, 52–58 (2019).

32. D. K. Jones, Diffusion MRI: theory, methods, and application (Oxford University Press, Oxford; New York, 2010), pp. xvi, 767 p.

33. J. A. Farrell et al., Effects of signal-to-noise ratio on the accuracy and reproducibility of diffusion tensor imaging-derived fractional anisotropy, mean diffusivity, and principal eigenvector measurements at 1.5 T. J Magn Reson Imaging 26, 756–767 (2007).

34. B. A. Landman et al., Effects of diffusion weighting schemes on the reproducibility of DTI-derived fractional anisotropy, mean diffusivity, and principal eigenvector measurements at 1.5T. Neuroimage 36, 1123–1138 (2007).

35. K. G. Schilling et al., Fiber tractography bundle segmentation depends on scanner effects, vendor effects, acquisition resolution, diffusion sampling scheme, diffusion sensitization, and bundle segmentation workflow. Neuroimage 242, 118451 (2021).

36. L. Ning et al., Cross-scanner and cross-protocol multi-shell diffusion MRI data harmonization: Algorithms and results. Neuroimage 221, 117128 (2020).

37. C. J. Molloy, S. Nugent, A. L. W. Bokde, Alterations in Diffusion Measures of White Matter Integrity Associated with Healthy Aging. J Gerontol A Biol Sci Med Sci 76, 945–954 (2021).

38. R. Nicolas et al., Changes Over Time of Diffusion MRI in the White Matter of Aging Brain, a Good Predictor of Verbal Recall. Front Aging Neurosci 12, 218 (2020).

39. C. A. Wheeler-Kingshott, M. Cercignani, About “axial” and “radial” diffusivities. Magn Reson Med 61, 1255–1260 (2009).

40. C. M. Tax et al., Cross-scanner and cross-protocol diffusion MRI data harmonisation: A benchmark database and evaluation of algorithms. Neuroimage 10.1016/j.neuroimage.2019.01.077 (2019).

41. D. K. Jones, P. J. Basser, “Squashing peanuts and smashing pumpkins”: how noise distorts diffusion-weighted MR data. Magn Reson Med 52, 979–993 (2004).

42. J. P. Fortin et al., Harmonization of multi-site diffusion tensor imaging data. Neuroimage 161, 149–170 (2017).

43. H. Mirzaalian et al., Inter-site and inter-scanner diffusion MRI data harmonization. Neuroimage 135, 311–323 (2016).

